# *Gli3* is required for glandular epithelial proliferation and endometrial homeostasis during the estrous cycle

**DOI:** 10.64898/2026.05.21.726971

**Authors:** Elizabeth Ung, Natalia M. Weinzierl, LeCaine J. Barker, Ashlyn N. Meinecke, Ryan M. Finnerty, Victor Ruthig, Elle C. Roberson

**Author notes:** Corresponding author information, Barbara Davis Center, CU Anschutz Medical Campus, 1775 Aurora Ct #A140, Aurora, CO 80045.

## Abstract

The endometrium is the innermost compartment of the uterus and undergoes cyclical remodeling throughout the human menstrual cycle and the rodent estrous cycle. The endometrium must thicken appropriately for embryonic implantation to occur; thus, it is crucial to understand the molecular mechanisms downstream of steroid hormone action that regulate endometrial thickness. Hedgehog (Hh) signaling is required for endometrial remodeling in both mice and humans, but the role of downstream Hh transcriptional effectors in endometrial remodeling is unknown. Here, we discover a role for the Hh transcriptional repressor, *Gli3*, in endometrial homeostasis: conditional knockout of *Gli3* resulted in a constitutively thick endometrium throughout the estrous cycle. In our model, a constitutively thick endometrium could support pregnancy. Bulk RNA-sequencing data revealed that loss of *Gli3* also resulted in dysregulated stromal-epithelial crosstalk, while immunofluorescent staining showed larger uterine glands and increased gland proliferation. These data deepen our understanding of molecular mechanisms controlling endometrial thickness, offering novel pathways to investigate endometrial factors in infertility.

## Introduction

The health of the non-pregnant, cycling endometrium is essential for fertility because receptivity to embryo implantation is dependent on endometrial thickness^1,2^. In humans, the endometrium rapidly regenerates and remodels throughout the menstrual cycle^3–5^. Beginning with menstruation, the upper two-thirds of the endometrium is shed, followed by endometrial cell proliferation and differentiation during the proliferative and secretory states, respectively^6^. The combination of proliferation and differentiation results in a thickened endometrium crucial for supporting implantation^7,8^. A thin endometrium is correlated with implantation failure, recurrent pregnancy loss, and later gestational complications like pre-eclampsia^1,2,8^. The mechanisms underlying appropriate endometrial thickening across the cycle, and how this confers later fertility are poorly understood. Here we report a crucial role of *Gli3* in managing endometrial thickness.

In the mouse and the majority of mammals, the endometrium does not shed via menstruation but endometrial thickening and thinning occurs during the analogous estrous cycle^9,10^. The endometrium responds cyclically to estrogen signaling, resulting in proliferation and thickening at proestrus and estrus, and then to progesterone signaling, resulting in cell death and endometrial thinning at metestrus and diestrus^9^. This occurs cyclically over a 5-7 day period in mice. Previously, we demonstrated that endometrial thickening is regulated by Hedgehog (HH) signaling, which is temporally activated at diestrus in the cycling mouse endometrium^10^. HH signaling was also shown to be necessary for implantation in myriad mouse models^11–14^. In humans, HH signaling is induced with similar timing during the menstrual cycle^15^, and decreased endometrial HH signaling is correlated with a thin endometrium and recurrent implantation failure^16,17^. Finally, in an ethanol-induced thin endometrium rat model, exogenous HH ligand restores endometrial thickness^18^.

The downstream effectors of HH signaling are the GLI transcription factors^19^. GLI3 is a transcriptional repressor required for mammalian limb and craniofacial development^20–22^, is implicated in human uterine development^23^, and is temporally upregulated at diestrus^10^. To test the role of *Gli3* in endometrial thickening, we conditionally knocked (cKO) out *Gli3*^24^ using the *progesterone receptor Cre* (*PR-Cre*)^25^, which is expressed in the uterine epithelium, stromal fibroblasts, and smooth muscle. The *Gli3* cKO produced a constitutively thick endometrium with dysregulated stromal-epithelial crosstalk and large, over-proliferative glands. Our findings suggest that HH signaling balances endometrial homeostasis through the cyclical actions of *Gli3*.

## Results

### Gli3 represses endometrial thinning

We assessed fertility and uterine homeostasis at estrus and diestrus in *Gli3* cKO, conditional heterozygotes (cHets), and controls (**Fig. 1A**). The partial or complete loss of uterine *Gli3* did not impact fertility at the level of live births per litter in our 3-month fertility trial (**Fig. 1B**). While fertility broadly describes many factors resulting in live birth, inspection of ovarian morphology revealed *Gli3* did not impact ovulation as noted by the unchanged number of corpora lutea in our model (**Supp. Fig 1A**). Further, systemic progesterone levels were not impacted by complete or partial loss of *Gli3* in tissues expressing *PR-Cre* (**Supp. Fig 1B**). Overall, with a normal birth rate, no changes in corpora lutea formation, nor progesterone levels, *Gli3* does not appear to significantly contribute to fertility in these tissues. While the importance of endometrial *Gli3* in fertility did not appear to be substantial, the morphologic differences in the endometrium when *Gli3* is lost were significant. Uterine, endometrial, and myometrial area expanded in estrus and shrank in diestrus when *Gli3* was expressed normally (**Figure 1D-1F**). When *Gli3* was lost (cKOs), or partially lost (cHets), the dynamic expansion and regression of the uterus, the endometrium, and the myometrium were lost (**Figure 1D-1F**). This was not due to changes in the myometrium or endometrium alone, as the ratio between these tissues did not significantly change in any condition (**Figure 1G**). This endometrial expansion was not associated with changes in all endometrial structures as the number of glands remained static across genotypes and cycle stages (**Fig. 1H**). We interpret these data to suggest that *Gli3* represses endometrial thinning, such that loss of *Gli3* results in a constitutively thick endometrium and subsequently normal fertility.

**Figure 1:**
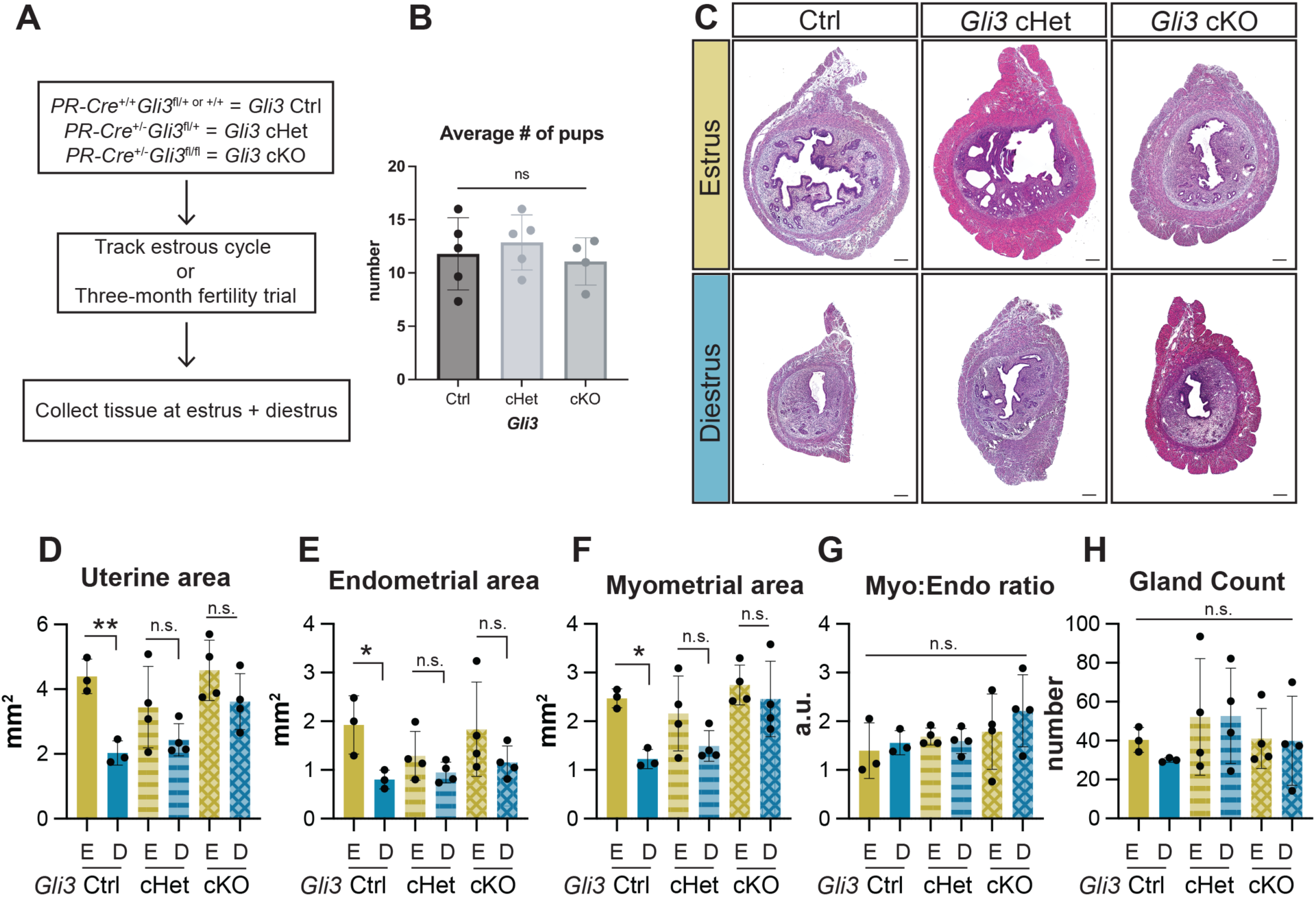
Gli3 represses endometrial thinning at diestrus. **(A)** Schematic of sample collection of Gli3 genetic littermate controls and mutants at estrus and diestrus. **(B)** The average number of pups from Gli3 controls, cHets, and cKOs, each dot represents the average number of pups for each biological replicate. **(C)** Hematoxylin and eosin staining of Gli3 control, cHet, and cKO uterine sections at estrus and diestrus. Scale bar = 200µm. Quantification of **(D)** total uterine area (**p-value=0.0032), **(E)** endometrial area (*p-value=0.0237), **(F)** myometrial area (*p-value=0.0113), **(G)** ratio of myometrial to endometrial area, and **(H)** gland count. Each dot represents the average of three or more sections across the uterine horn of a biological replicate. Control estrus and diestrus n=3, cHet estrus and diestrus n=4, cKO estrus and diestrus n=4. A two-way ANOVA with multiple comparisons of cell means with others in its row and column was run for statistical significance. n.s. = not significant.

### Hedgehog signaling is dysregulated in the absence of Gli3

To understand how loss of endometrial *Gli3* affected HH signaling, we assessed gene expression of HH signaling components including the endometrial ligand (*Ihh*), the canonical gene targets (*Ptch1, Gli1*), the signaling effector (*Smo*), and the transcription factors (*Gli2*, *Gli3*) (**Fig. 2**). *Ihh* is a progesterone target gene^14,26^ that we previously showed as upregulated at diestrus in control uteri^10^. *Gli3* controls, cHets, and cKOs upregulated *Ihh* expression during diestrus, as expected (**Fig. 2A**), suggesting that *Gli3* does not regulate upstream ligand expression. Downstream canonical targets, *Ptch1* and *Gli1*, fail to significantly increase gene expression during diestrus in *Gli3* cKO compared to control uteri (**Fig. 2B, C**) suggesting that HH signaling is dysregulated in the *Gli3* cKO uteri. We further assessed *Smo* and *Gli2* expression and found no differences across genotypes (**Fig. 2D, E**), similar to *Ihh* expression patterns. Finally, we validated that *Gli3* expression is significantly decreased or lost in *Gli3* cKO uteri across estrous stages (**Fig. 2F**).

**Figure 2:**
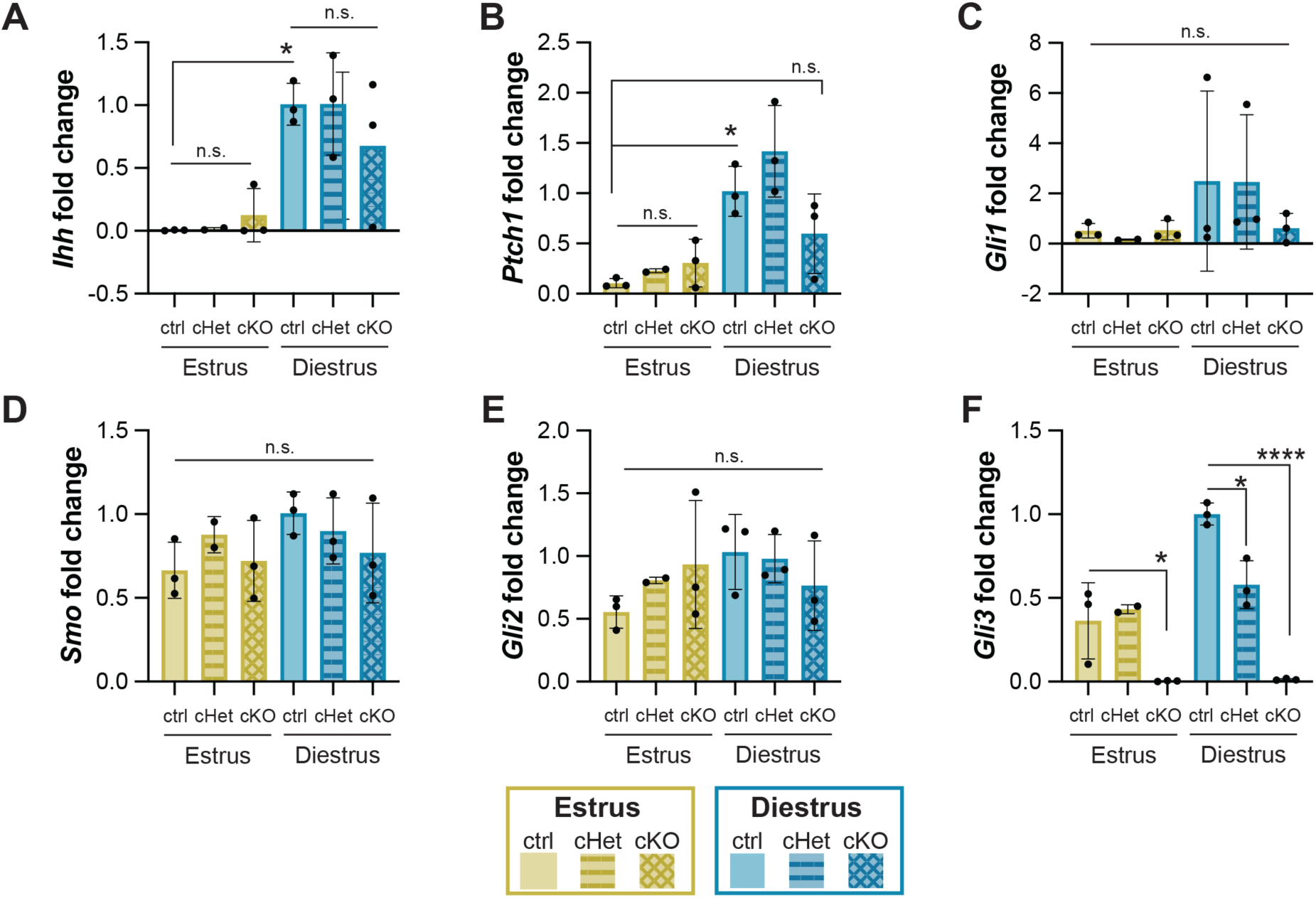
Hedgehog signaling is dysregulated in the absence of Gli3. (A-F) qPCR quantification of Hh signaling components, Ihh (A), Ptch1 (B), Smo (D), Gli1 (C), Gli2 (E), and Gli3 (F) at estrus and diestrus in whole uterus lysate of control, cHet, and cKO. Each dot represents one biological replicate, which is the average of four technical replicates. *p-value < 0.05; ****p-value <0.00001; n.s. = not significant.

### Gli3 regulates stromal-epithelial crosstalk

At implantation, epithelial-derived IHH is a significant regulator of epithelial-stromal crosstalk^11,14,27^, but HH gene targets in the cycling uterus are unknown. To define how *Gli3* contributes to epithelial-stromal crosstalk, we performed bulk RNA sequencing (RNA-seq) to identify differentially regulated genes in *Gli3* controls and cKOs at estrus and diestrus. In control samples, 1157 genes were upregulated at estrus compared to diestrus, and 989 genes were downregulated (**Fig. 3A**). To determine within-stage how *Gli3* regulates gene expression, we asked how the cKO compares to controls at each stage. At estrus, the *Gli3* cKOs displayed aberrant gene expression with 231 upregulated genes and 228 downregulated genes (**Fig. 3B**). *Gli3* also regulated gene expression at diestrus, with 252 and 308 upregulated and downregulated genes, respectively (**Fig. 3C**). In control animals, we saw similar cycling of gene expression between estrus and diestrus as previously reported^10^ (**Fig. 3D**). At both estrus and diestrus, the *Gli3* cKO displayed de-repressed and inactivated genes (**Fig. 3E, F**). To identify biological pathways dysregulated by loss of *Gli3*, we performed GO term analysis^28,29^. In controls, we found that metabolic functions, response to estradiol, and negative regulation of cell adhesion were upregulated at estrus (**Supp.Fig. 2A**) while functions relating to synapse and neural activity were upregulated at diestrus (**Supp. Fig. 2B**). In *Gli3* cKOs at estrus, we found that “immune response” was upregulated (**Supp. Fig. 2C**), suggesting that *Gli3* may repress immunological or inflammatory responses. In *Gli3* cKOs at diestrus, we found downregulation of GO terms related to postsynaptic activity (**Supp. Fig. 2D**). Intriguingly, morphogenesis of branching epithelium, regionalization, and artery and lung development were a few outstanding GO terms that were downregulated in the cKO at estrus (**Fig. 3G**, bolded), suggesting that stromal *Gli3* regulates epithelial biology. In *Gli3* cKOs at diestrus, we found upregulation of proliferation associated GO terms like DNA replication, cell cycle, cholesterol metabolic process, and fibroblast proliferation as well as regulation of tube diameter (**Fig. 3I**, bolded and italicized, respectively) suggesting that *Gli3* may repress cellular proliferation and regulate tube size. Uterine glands are relevant epithelial tubes located in the endometrium that undergo cyclical morphogenesis (remodeling and regeneration) across the estrous cycle^30–32^. More detailed heatmaps of representative genes from relevant GO terms, including morphogenesis of branching epithelium and proliferation are provided in the supplemental data (**Supp. Fig. 3**). Our gene expression dataset suggests that *Gli3* coordinates stromal-epithelial crosstalk by regulating gland morphogenesis gene expression at estrus and robustly repressing cellular proliferation genes at diestrus.

**Figure 3:**
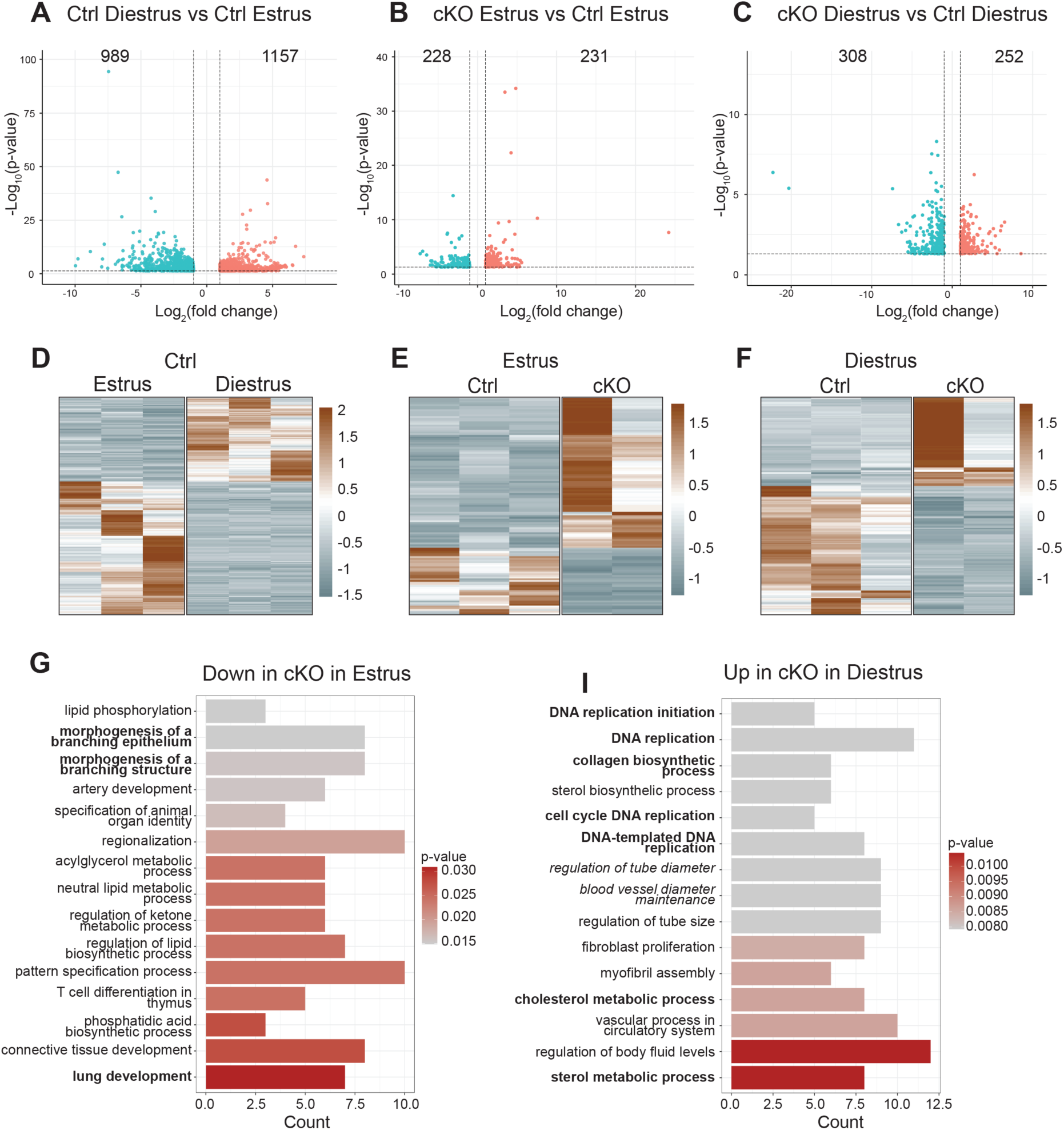
Gli3 regulates stromal-epithelial crosstalk. (A-C) Volcano plots showing the number of up and downregulated genes at estrus compared to diestrus in Gli3 controls (A), and cKOs animals compared to controls at estrus (B) and diestrus (C). **(D-F)** Heatmaps showing the most significant (p-value < 0.01) differentially expressed genes in Gli3 controls (D) and cKOs animals at estrus (E) and diestrus (F). **(G-I)** GO term analysis of downregulated genes in the cKOs at estrus (G) and upregulated genes in the cKOs at diestrus (I).

### Gli3 maintains gland size

Our bulk RNA-seq dataset prompted us to take a closer look at uterine glands as we observed transcriptional evidence of decreased gland branching at estrus, suggesting that the *Gli3* cKO was associated with a gland phenotype. We had anecdotally noted some large glands in the *Gli3* cKO in our original H&E staining (**Fig. 1C**), so we identified how many uteri within our dataset displayed at least one large gland (**Fig. 4A**). A higher percentage of *Gli3* cHet and cKO uteri displayed larger glands (**Fig. 4B**). When we assessed when large glands appeared across the estrous cycle, *Gli3* cHets and cKOs had large glands throughout the cycle, whereas the control animal with large glands appeared only at estrus (**Fig. 4C**). While the *Gli3* cHet and cKO uteri had larger glands, the gland epithelium was organized and cuboidal as expected (**Fig. 4A**), and we found no evidence of hyperplastic glands like that seen in precursors to endometrial cancer^33,34^. We assessed gland size quantitatively and found that the *Gli3* cHet and cKO animals had a higher percentage of ‘large’ glands compared to their control counterparts (**Fig. 4D**). When we specifically plotted the number of extremely large glands (>1000µm^2^), we saw a significant increase in the number of glands between control, cHet, and cKO uteri at estrus (**Fig. 4E**). We confirmed that uterine glands were specified appropriately by visualizing the transcription factor Forkhead box a2 (FOXA2)^35–37^ and E-CADHERIN (E-CAD) (**Fig. 4F**). Finally, we visualized an entire *Gli3* cKO uterine horn to confirm that larger glands appear throughout the entire horn (**Supp. Fig. 4**).

**Figure 4:**
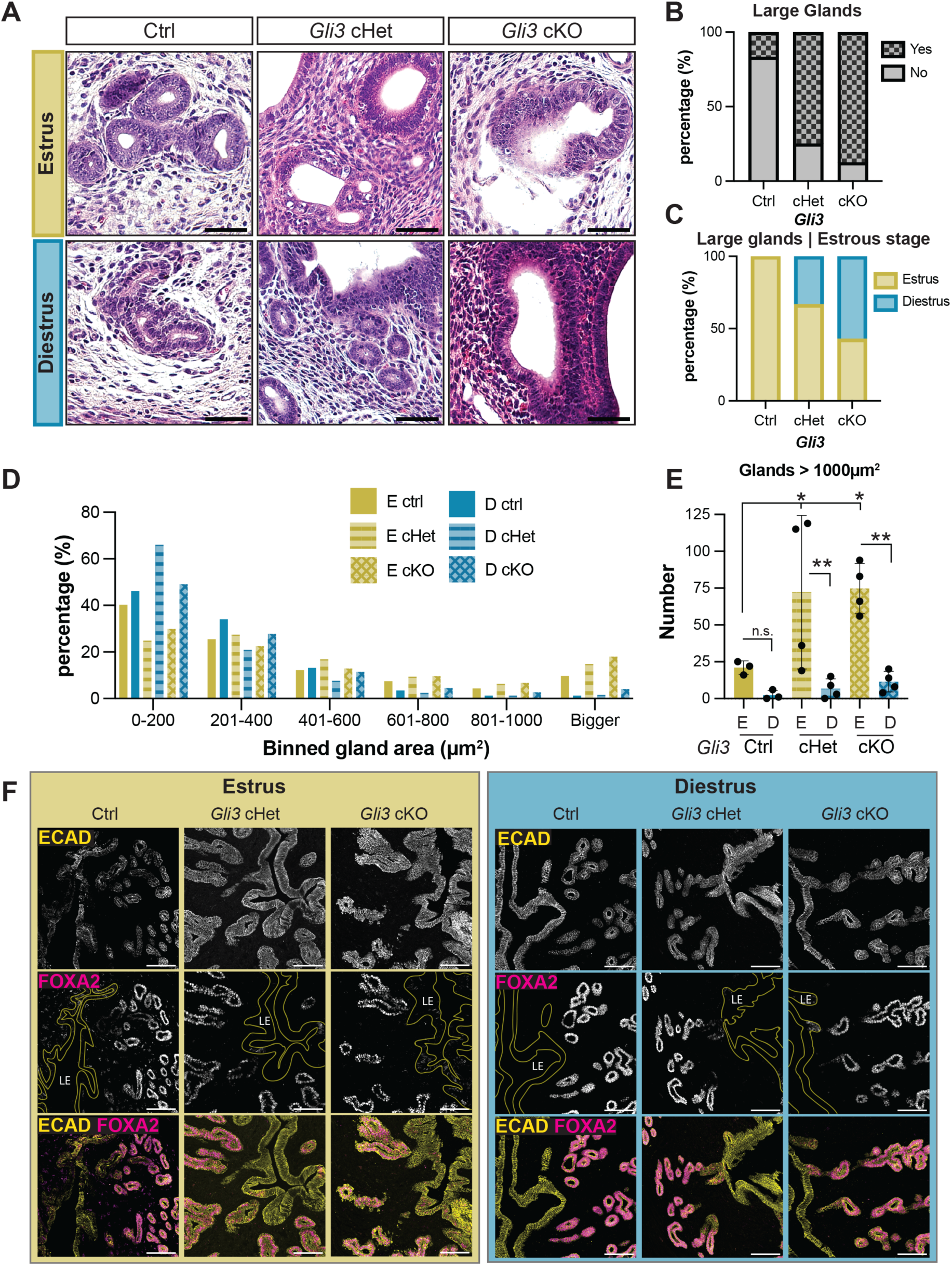
Gli3 maintains gland size. **(A)** Hematoxylin and eosin staining of Gli3 littermate controls, cHets, and cKOs, focusing on uterine glands. Imaged with a 20X objective on a Lecia DM6b brightfield microscope. Scale bars = 50µm. **(B)** Quantitation of the percent of animals displaying large glands. **(C)** Quantitation of the percent of animals with large glands in each estrous cycle stage. Quantitation of **(D)** the percentage of glands per binned area, and **(E)** the total number of glands that are larger than 1000µm^2^ (**p-value < 0.0020, *p-value <0.0402). Each dot represents the average area (µm^2^) of glands in one biological replicate. Control estrus and diestrus n=3, cHet estrus and diestrus n=4, cKO estrus and diestrus n=4. A two-way ANOVA with multiple comparisons of cell means with others in its row and column was run for statistical significance. **(F)** Immunofluorescent staining of Gli3 control, cHet, and cKO uterine sections with FOXA2 (magenta) and E-CAD (yellow) at estrus (A) and diestrus (B). LE = luminal epithelium. Scale bars = 100µm.

### Gli3 represses gland epithelium proliferation

Differences in gland size could arise from changes in epithelial proliferation or apoptosis. Because our bulk RNA-seq dataset suggested that *Gli3* represses proliferation genes at diestrus, we hypothesized that the loss of *Gli3* would increase gland epithelial proliferation at diestrus. To test this hypothesis, we labeled all proliferating cells with KI67 and all epithelium with E-CAD (**Fig. 5A**). We noted increased KI67 staining in gland epithelium at diestrus in our *Gli3* cHets and cKOs, as predicted. Therefore, we quantified proliferation of glandular epithelium across genotypes and estrous cycle stages using Imaris (**Fig. 5B, Supp. Fig. 5**). From our z-stacks of E-CAD, we segmented the 3D surfaces of both luminal and glandular epithelium (**Fig. 5B, Supp. Fig. 5**). We used a size-based filter to remove the luminal epithelium from analysis and then manually validated this filtering to remove any remaining luminal epithelial artifacts. Within the identified gland epithelium region of interest, we segmented the 3D surface of proliferative cells using KI67 (**Fig. 5B, Supp. Fig. 5**). We then calculated the percent volume of proliferative cells within the glandular epithelium (**Fig. 5C**). In our control animals, we found no difference in GE proliferation (**Fig. 5C**), likely because GE proliferation occurs during proestrus and metestrus^9^. When we assessed GE proliferation across genotypes at estrus, we found no significant differences (**Fig. 5C**). However, when we assessed GE proliferation across genotypes at diestrus, we found a significant increase in the cHets and a trending increase in the cKOs (**Fig. 5C**). Haploinsufficiency of *Gli3* is known to produce phenotypes, like polydactyly^38^, so we were not surprised to find that the *Gli3* cHet displayed phenotypes similar to the cKO animals. These data suggest that glands are larger in *Gli3* cHets and cKOs due to aberrantly increased proliferation at diestrus.

**Figure 5:**
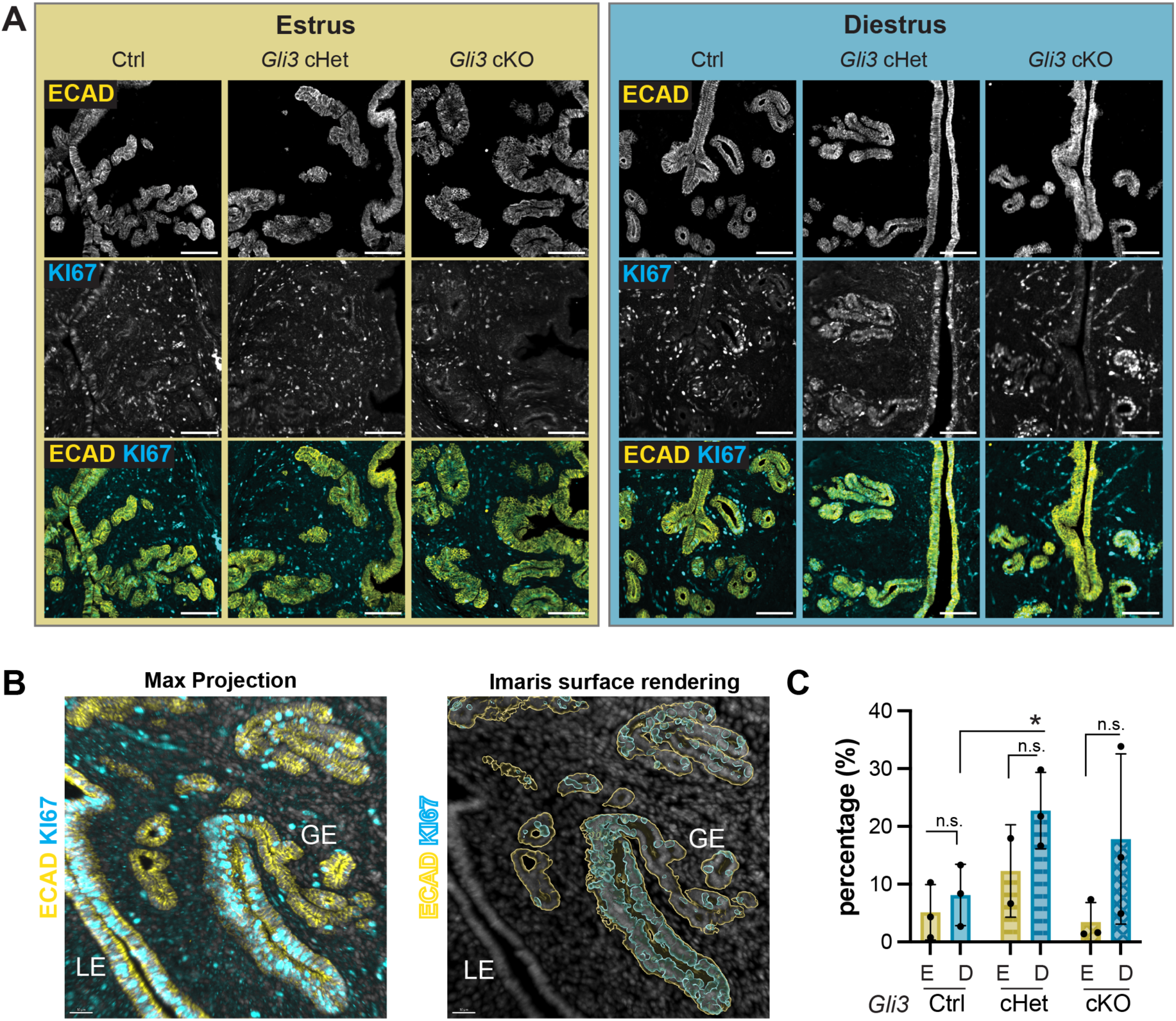
Gland epithelium is more proliferative in Gli3 cKOs. **(A)** Immunofluorescent staining of Gli3 controls, cHets, and cKOs for E-CADHERIN (yellow, pan-epithelial marker) and KI67 (cyan, proliferation marker) at estrus and diestrus. Scale bar = 100µm. **(B)** Representative Imaris image to surface render gland epithelium and KI67, used to quantify KI67 volume within gland epithelium volume. **(C)** Quantification of KI67 volume within gland epithelium volume, plotted as percentage. Each dot represents one biological replicate with an average of at least 3 technical replicates. *p-value < 0.05; n.s.= not significant.

## Discussion

Endometrial thickness is an important indicator of fertility; a thin endometrium is correlated with recurrent pregnancy loss^1,2^. Previous work has shown that HH signaling is required for fertility and endometrial remodeling^10,11,27,39,40^, however, the molecular mechanisms of how HH signaling regulates endometrial thickening remain poorly understood. Here, we provide evidence that the HH transcription factor, *Gli3*, regulates endometrial thickness through stromal-epithelial crosstalk controlling epithelial proliferation and gland size across the estrous cycle.

The uterine HH ligand *Ihh* is expressed by luminal epithelium and signals to the stromal fibroblasts in order to regulate epithelial-stromal crosstalk and stromal proliferation at implantation^11,27,39,41^. Our work significantly builds on this by integrating *Gli3* into the mechanistic model of endometrial HH signaling. Our experiments revealed both upregulated (de-repressed) and downregulated (inactivated) gene expression in *Gli3* cKOs compared to controls at both estrus and diestrus. While GLI3 acts primarily as a strong transcriptional repressor, it also has weak activator activity^42^. In the absence of HH ligand, GLI3 is proteolytically processed into a repressor^19^, suggesting a model whereby GLI3 represses downstream HH target genes in the absence of signaling^43^. In the uterus, HH signaling is activated temporally at diestrus^10^, suggesting temporary loss of GLI3 repressor and thus de-repression of GLI3 target genes. However, the timing of GLI3 repression is tissue dependent. For example, during limb bud development, GLI3 does not repress HH target genes until after HH signaling occurs^44^. While we speculate that GLI3 is repressing genes at other estrous cycle stages, further experimental analysis is required to identify how GLI3 is behaving as a transcription factor in the endometrium.

In the *Gli3* cKOs, inflammatory GO terms were upregulated at estrus, suggesting that GLI3 represses inflammation at this stage. In line with this observation, in the wild-type cycling uterus inflammation-associated genes are upregulated at estrus^45^, and the uterus heals more rapidly and efficiently when injured at estrus compared to diestrus^46^. The stage of uterine injury had a significant effect on pregnancy: dams injured at estrus had normal spacing and placentas, whereas diestrus injury resulted in increased embryo death, abnormal spacing, and placenta accreta-like phenotypes^46^. While there are several possible explanations for this, inflammation plays a clear role in repair and regeneration^47^, suggesting that estrus is primed for regenerative capacity. GLI3 may play a role in balancing endometrial inflammation by repressing relevant genes. Aberrant inflammatory mechanisms are associated with endometrial disease, like endometriosis, driving pathological proliferation and tissue adhesions^48^. GLI3 and other HH signaling components are upregulated in endometriosis lesions^49^, suggesting a potential connection between HH-associated inflammatory regulation and endometriosis progression. There is precedent for this in other diseases, as HH signaling is associated with inflammation in other organs like the small intestine, where reduced signaling is implicit in inflammatory bowel disease^50^.

We also found that loss of *Gli3* results in dysregulated stromal-epithelial crosstalk at the transcriptional level: in *Gli3* cKOs, branched epithelium GO terms are downregulated at estrus, and proliferation GO terms are upregulated at diestrus. Uterine glands undergo initial branching morphogenesis during development^51,52^ and further branching and coiling in early pregnancy to support implantation^53,54^. However, how glands balance proliferation and branching is poorly understood. Loss of *Gli3* resulted in an increase in glandular epithelium proliferation in diestrus, suggesting that GLI3 represses gland proliferation and weakly (or indirectly) activates gland branching. We hypothesize a weak or indirect activation of gland branching because branching is required for gland function and fertility^54^. Since *Gli3* cKO mice are fertile, this suggests that gland branching occurs normally. However, this balance of pro-proliferative and anti-branching may explain why we observe larger glands throughout the estrous cycle. Interestingly, in the mammary gland, stromal cells are proposed to control mammary gland epithelial proliferation, whereas mammary gland morphogenesis is regulated intrinsically by the epithelium^55^. We propose a similar model in the uterus, where stromal *Gli3* regulates gland proliferation, but gland branching is likely epithelial-intrinsic.

In sum, we propose a model wherein GLI3 plays a repressive regulatory role in maintaining appropriate endometrial thickness by regulating proliferation genes and stromal-epithelial crosstalk across the estrous cycle. Canonically, pathway activation prevents the accumulation of GLI3 repressor (GLI3-R), leading us to an intriguing model of cyclically regulated GLI3 function. Given that HH signaling is transiently upregulated at diestrus^10^, we suggest that GLI3-R is temporarily lost at diestrus resulting in de-repressed expression of genes involved in proliferation. At subsequent cycle stages (proestrus, estrus, and metestrus), GLI3-R is restored, thus repressing over-proliferation and inflammation-associated genes, while allowing for weak activation of epithelial branching genes. Future work will continue to test this model. It is crucial to understand the molecular mechanisms controlling endometrial thickness, as recurrent pregnancy loss can be due to a thin uterus. These data contribute to our understanding of endometrial homeostasis, and therefore how individuals may experience infertility.

### Limitations of the study

Analyzing the estrus and diestrus stages allows for a broad comparison where overall tissue, cellular, and molecular level changes are distinct. However, we are limited because gland proliferation occurs at proestrus and metestrus^30^, thus our estrous stage sampling is not ideal to validate proliferation in the control animals. Nonetheless, we identified aberrant proliferation at diestrus in the *Gli3* cHets and cKOs. Sampling additional estrous stages would strengthen these data but was outside the scope of this manuscript. Performing bulk RNA-sequencing also limits our ability to study the different cell types within the endometrium, including but not limited to, the luminal and glandular epithelium, and the stromal cells surrounding them. Finally, we used *PR-Cre* that is expressed as early as at postnatal day 5 in most cells of the uterus as well as other progesterone-responsive tissues. While we predict that our phenotypes are from loss of stromal *Gli3,* we are unable to rule out that our phenotypes arise from loss of *Gli3* in other cell types within the uterus.

## Acknowledgements

We thank the CU Anschutz vivarium and Office of Laboratory Animal Research (OLAR) for their work in supporting our mouse colony. We would like to thank the University of Colorado Denver Histology Shared Resource Core and the University of Virginia Center for Research in Reproduction Ligand Assay and Analysis Core. We would like to thank all members of the Roberson lab for critical discussion of the work presented. We would also like to thank Dr. David Haimes. This work was supported by start-up funds to E.C.R. provided by CU Anschutz School of Medicine.

## Author Contributions

**E.U.:** Investigation, Methodology, Validation, Data Curation, Formal Analysis, Bioinformatic Analysis, Writing. **N.M.W.:** Investigation, Validation. **L.J.B.:** Methodology, Investigation, Validation. **A.N.S.:** Bioinformatic Analysis. **R.M.F.:** Investigation, Validation. **V.R.:** Image Analysis. **E.C.R.:** Conceptualization, Methodology, Validation, Formal Analysis, Writing, Supervision, Project Administration, Funding Acquisition.

## Materials and Methods

### Mice

Mice were housed in individually ventilated cages in a pathogen-free facility with continuous food and water. Homozygous floxed *Gli3* females (*Gli3^fl/fl^*, JAX: 008873;^24^) and heterozygous PR-Cre males (*PR-Cre^+/-^,* JAX: 017915;^25^) were bred over two generations to produce experimental (*PR-Cre^+/-^;Gli3^fl/fl^* or *PR-Cre^+/-^;Gli3^+/-^*) or control (*PR-Cre^+/+^;Gli3^fl/+^* or *PR-Cre^+/+^;Gli3^fl/fl^*) females. Animals were housed with a 12hr on/off light cycle in individually ventilated cages. Mice were humanely euthanized via isoflurane followed by cervical dislocation, and the entire female reproductive tract was dissected. All experiments were approved by the University of Colorado Anschutz Institutional Animal Care and Use Committee (IACUC #01267).

### Estrous cycle tracking

8-22-week-old females were estrous cycle checked using vaginal cytology^56^. Animals were first acclimated to estrous cycle tracking for a week by briefly swabbing the vaginal canal using a cotton swab that was wetted with 1X PBS. After a week of acclimation, the animals were tracked through the same method, except the swabs were smeared onto a microscope slide. Slides were stained with crystal violet (Fisher Scientific, cat# C581-25), and the estrous stage was determined using a brightfield microscope (Leica DM6b). This protocol persisted until the animal was in the correct estrous stage for dissections.

### Fertility trials

Up to two female mice were placed into a single male cage by 4 p.m. to mate. Females were plug checked every morning thereafter using a vaginal probe until a plug was identified. If a female was found with a plug, she was removed from the male cage and housed separately. Plugged females were checked daily for the delivery of pups starting at embryonic day (E) 17.5. Once delivered, pups were euthanized through decapitation. The dam was housed alone for five days before being placed back into mating. Tracking persisted up to 3 sequential litters or three months.

### Tissue processing

#### Dissections

After euthanasia and cervical dislocation, cardiac puncture was completed to collect blood to assess serum progesterone levels analysis. The entire reproductive tract of each animal was dissected out, and the uterus was bisected at the cervix, utilizing one uterine horn for paraffin embedding and processing, and RNA collection, and the other horn for cryopreservation and whole mount tissue processing. Each uterine horn was laid on a strip of index card to ensure proper structure of the uterine horns during fixation. For RNA, uterine tissue was stored at-20°C in RNA*later* (Invitrogen, cat# AM7021).

#### Fixation

Samples were fixed in 4% paraformaldehyde/1X PBS (Gibco, cat# 70011-044) for 3 hours at 4°C. After fixation, samples were washed in 1X PBS 3×5 minutes while rocking at room temperature. Samples were then put through proper post-processing. Uterine horns sent for paraffin processing were placed in 70% ethanol overnight at room temperature before paraffin embedding at the University of Colorado Denver Research Histology Shared Resource Core (RRID: SCR_021994). Uterine horns used for cryopreservation were placed into 30% sucrose (in 1X PBS) overnight at 4°C. Uterine horns used for whole mount tissue clearing and staining were incubated in an isopropanol/1X PBS gradient of 30%, 50%, 70%, and 100%, for 20 minutes at each concentration^57^. Samples were then stored at-20°C in 100% isopropanol until ready for use.

#### Cryopreservation embedding

Samples were taken out of 30% sucrose and blotted dry on a kimwipe. A 15 × 15 × 5 mm cryo mold was filled with OCT (Fisher Scientific, cat# 23720571) before uterine horns were cut into 0.5 cm sections and placed transversely into the mold. The mold was placed into a dry ice and 100% ethanol bath to gradually freeze. Tissue blocks were stored at-80°C until ready to section.

#### Serum analysis

Collected blood from estrous checked animals were placed in a 1.5mL microcentrifuge tube and set to clot at room temperature for 60-90 minutes. Following clotting, samples were centrifuged for 15 minutes at 2000 x g. Serum was collected and placed in a new 1.5mL microcentrifuge tube and immediately stored at-80°C before being sent out to the University of Virginia Center for Research in Reproduction Ligand Assay and Analysis Core for progesterone hormone analysis.

#### Tissue sectioning

#### Paraffin

Paraffin blocks were placed in an ice/water bath to cool down the paraffin wax before being placed into a Leica RM2155 microtome. The block was exposed, and tissue was transversely cut at 5µm. Ribbons were placed into a warm water bath and maneuvered onto Superfrost plus microscope slides (Fisherbrand, cat# 1255015). Slides were dried overnight at room temperature before storing in slide boxes indefinitely at room temperature.

#### Cryo

Cryopreserved blocks adjusted from-80°C to-20°C for 30-60 minutes before mounting on a cold chuck with OCT. The block was exposed and sections collected at 12µm across the entire uterine horn on Superfrost plus microscope slides (Fisherbrand, cat# 1255015). Slides were stored long term at-80°C.

### Histology and quantitation

#### Hematoxylin and eosin (H&E) staining

The StatLab H&E stain kit (cat# KTHNEPT) was used. Paraffin sectioned slides were baked at 60°C for 20 minutes. The slides were deparaffinized by immersing slides 2×5 minutes in xylene. Slides were then dehydrated with 100% ethanol 3×1 minutes. Slides were rinsed in running tap water for 1 minute before being immersed in hematoxylin for 5 minutes, followed by another rinse in running tap water for 1 minute. The slides were then dipped 2-3 times for 1 second each in differentiating solution, rinsed in running tap water for 1 minute, and immersed in bluing solution for 30-40 seconds before a final rinse in running tap water. Samples were placed in 70% ethanol for 1 minute before incubating with Eosin Y for 30-60 seconds, followed by a 95% ethanol incubation for 30-45 seconds. Slides were dehydrated with 100% ethanol for 3×1 minutes followed by 3×1 minute xylene incubation. Samples were coverslipped and cured using Cytoseal 60 (Epredia, cat# 23-244257) overnight before imaging on a Leica DM6b brightfield microscope using 10x (Leica, cat# 506410) and 20x objectives (Leica, cat# 506529) as color images and exported as.tif files.

#### Areas

H&E-stained tissue images were imported into FIJI (version 2.14.0) to measure total uterine area, endometrium area, and gland count^58^. *Total uterine area*: raw tif file images were converted to 8-bit and thresholded before using the wand tracing tool to select the entire uterus to measure. *Endometrium area*: using the polygon selection tool, the endometrium/circular smooth muscle layer border was traced and measured. *Gland count*: the multipoint tool was used to count each gland in the uterine section. *Myometrium area*: generated by subtracting endometrium area from total uterine area. *Myo:endo area ratio*: generated by dividing myometrium area by endometrium area. Three sections distributed evenly across the uterine horn were measured. Uterine area, endometrium area, and gland count were averaged across sections for each animal and graphed in GraphPad Prism (version 11.0.0(93)), where a two-way ANOVA test across multiple comparisons was run for statistical analysis.

#### Gland size

H&E-stained tissue images were placed into QuPath (version 0.5.1)^59^. Using the wand tool, individual gland lumen area (μm^2^) was measured, excluding the glandular epithelium. Glands from four sections distributed evenly across the uterine horn were measured. Average gland size was calculated and sorted into 200, 400, 600, 800, 1000μm^2^, or bigger bins. A two-way ANOVA test across multiple comparisons was run for statistical analysis in GraphPad Prism.

#### Immunofluorescence, imaging, and quantitation

*Immunofluorescence staining*. Cryosectioned slides were set out to adjust to room temperature for 15-20 minutes before washing OCT off with 3×5 minute 1X PBS washes. Samples were blocked with blocking solution made of 10% TritonX-100 (Thermo Fisher Scientific, cat# NC2034995), 5% normal donkey serum (Jackson Immuno Research, cat# NC9624464) and 1X PBS for 30-60 minutes at room temperature. The following primary antibodies were used and incubated overnight at 4°C in a humidity chamber: anti-KI67 (rabbit, 1:1000, Abcam, AB15580) and anti-E-CADHERIN (goat, 1:500, R&D Systems, AF748), anti-FOXA2 (rabbit, 1:250, Abcam, AB108422). Samples were washed 3×5 minutes with PBS. The following secondary antibodies were used and incubated for 30 minutes-2 hours at room temperature: AlexaFluor anti-rabbit 647 (1:1000, Invitrogen, A31573), AlexaFluor anti-goat 488 (1:1000, Invitrogen, A11055), and DAPI (1:1000, Millipore Sigma, 5087410001) (**Table 1**). Slides were washed 3×5 minutes with 1X PBS before coverslipping with Prolong Diamond Antifade Mountant (Invitrogen, P36970). Images were collected on an Andor Dragonfly 200 spinning disk confocal microscope (Oxford Instruments) using Fusion software (version 2.4.0.14) with a 10x (NA 0.45) air objective (Leica, cat# 11506410) with 405nm, 488nm, and 561nm or 638nm diode lasers for excitation, and a piezo stage (ASI) was used to acquire Z stacks at 5µm intervals. Image pre-processing was done in Fusion (v2.4) to stitch tile-scanned (montaged) images.

**Table 1.** Primary and secondary antibodies.

| Antibody | Host | Company | Catalog # | Dilution |
| --- | --- | --- | --- | --- |
| KI67 | Rabbit | Abcam | AB15580 | 1:1000 |
| E-CADHERIN | Goat | R&D Systems | AF748 | 1:500 |
| FOXA2 | Rabbit | Abcam | AB108422 | 1:1000 |
| FOXA2 | Rabbit | Cell Signaling | 8186S | 1:200 |
| AlexaFluor anti-goat 488 secondary | Donkey | Invitrogen | A11055 | 1:1000 |
| AlexaFluor anti-rabbit 647 secondary | Donkey | Invitrogen | A31573 | 1:1000 |
| DAPI | N/A | Millipore Sigma | 5087410001 | 1:1000 |

#### Cell proliferation quantitation

Image analysis was performed in Imaris (version 11.0.1, Oxford Instruments Bitplane). Samples were labeled only with stage of the estrous cycle so that analysis was blinded to genotype. An analysis pipeline was built within the Imaris Workflow environment. Four estrus and four diestrus samples were randomly selected to build and initially validate a segmentation Workflow. The initially validated Workflow was then tested on the full set of samples for secondary validation to develop a final Workflow. The final Workflow was then applied to all the samples and the data collected for analysis and images collected for figure making. The Workflow file is available upon request.

The Workflow proceeded as follows. Pre-processed anti-E-CADHERIN stain was used to segment glandular epithelium with the Surfaces model. The absolute intensity algorithm was used for initial segmentation which was further refined with a filter for the number of voxels to distinguish between glandular epithelium segmentations which were kept, and uterine epithelium segmentations which were removed. Occasionally sheared parts of uterine epithelium or autofluourescent uterine debris passed the filtering step. These were independently verified as “not glandular epithelium” and then removed from the segmentation as part of a manual intervention and manual cleaning step in the Workflow. Preprocessed anti-KI67 stain was used to segment active cells with the Surfaces model. The region of interest (ROI) for the Surfaces model was constrained to only the glandular epithelium area defined previously with Surface segmentation on anti-E-CADHERIN stain. Within this ROI, the absolute intensity algorithm was used for initial segmentation of anti-KI67 stain which was further refined with a filter for the number of voxels to remove noise too small to be real signal. Raw data tables exported from the Imaris Workflow for each sample included data for volumes of glandular epithelium and active cells. An analysis pipeline in R (version 4.5.2)^60^ was executed in Rstudio (version 2026.01.1+403)^61^ using packages tidyverse^62^, readxl^63^, writexl^64^, openxlsx^65^, purr^66^, dplyr^67^, plyr^68^, and tidyr (**Table 2**). The analysis pipeline read in the paired volumes for each sample, aggregated all the data into a new single table, and calculated percent active values for glandular epithelium. The percent active data was imported into GraphPad Prism (version 11.0.0) for graphing and statistical analysis. Maximum intensity projections of views of the sample image exported from the Imaris Workflow were used for example figures which were made in Adobe Illustrator (version 30.3).

**Table 2.**
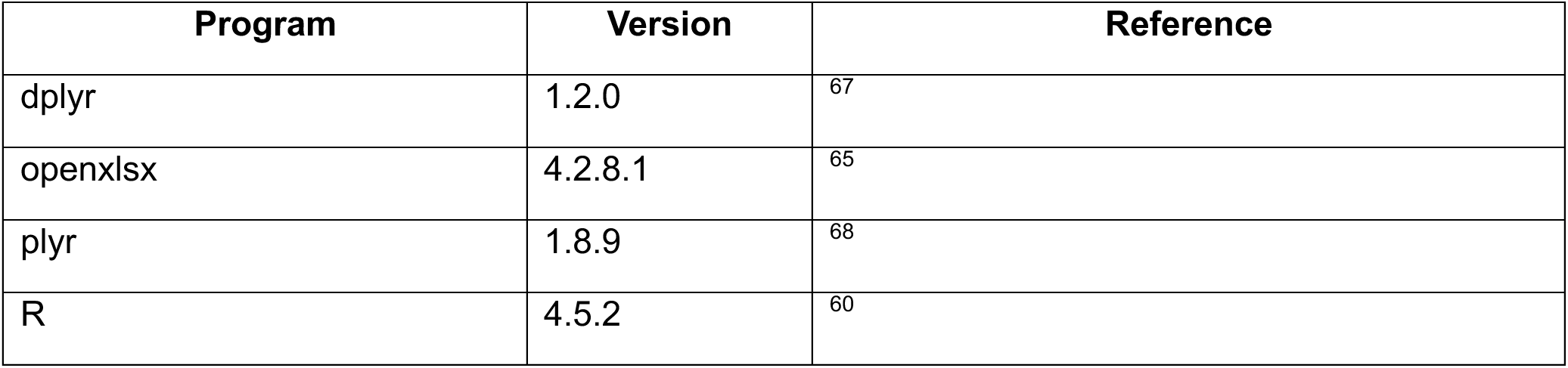
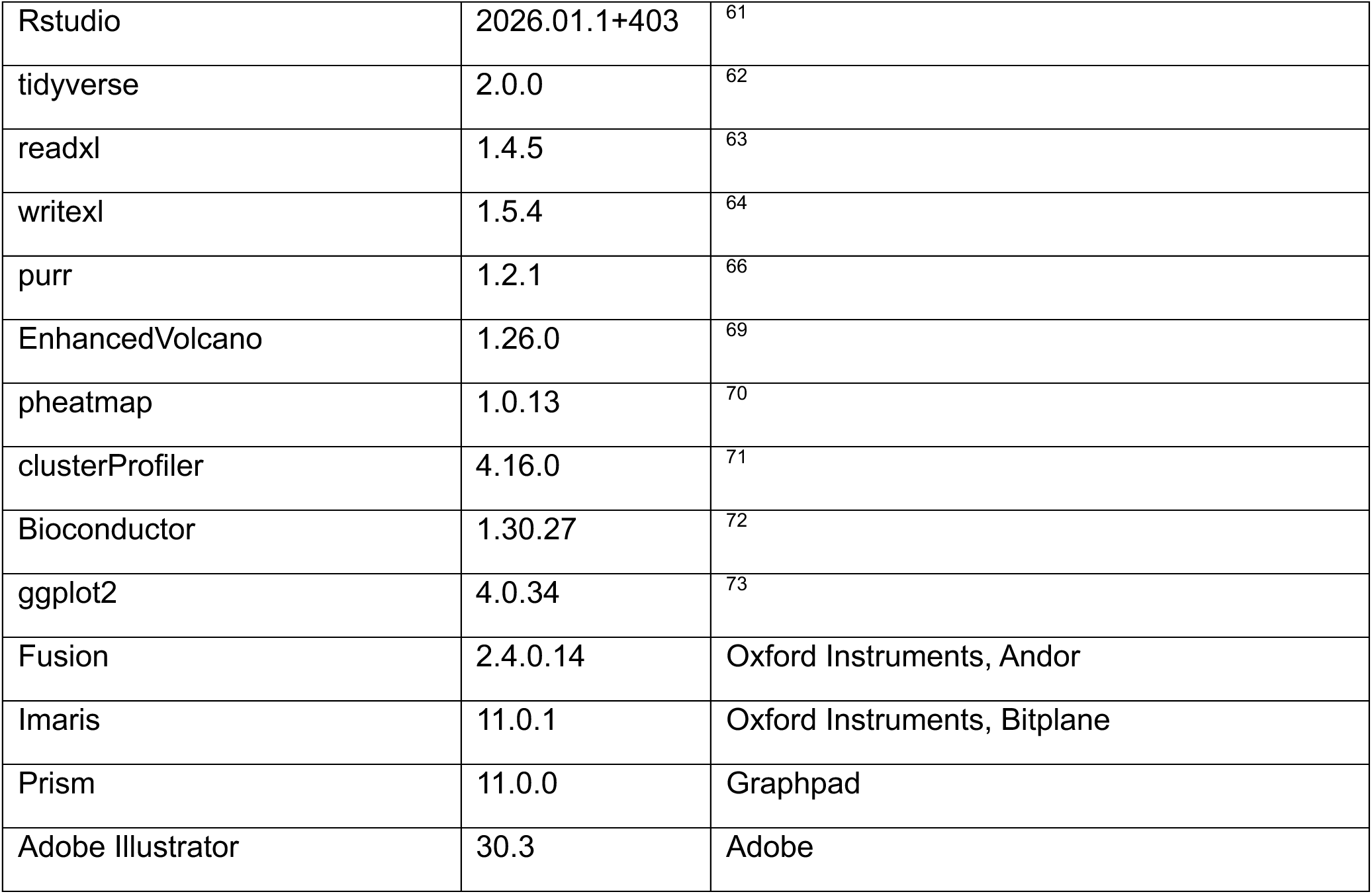

#### Whole mount tissue clearing

Whole mount tissue clearing was performed as previously published^57^. Briefly, uterine horns were rehydrated from 100% isopropanol storage with isopropanol/1X PBS gradient washes from 70%, 50%, and 30%. Samples were then permeabilized overnight at 37°C. The next day, samples were blocked for 5-6 hours at 37°C before placing in antibody solution with rabbit anti-FOXA2 (rabbit, 1:200, Cell Signaling, 8186S) and anti-E-CADHERIN (goat, 1:500 dilution, R&D Systems, AF748) for three nights at 37°C. After primary antibody incubation, samples were washed 3×1 hour in PBS/TritonX-100 with heparin and incubated with AlexaFluor anti-rabbit 647 (Jackson Immuno Research, NC0254454) and Cy3-Affinipure anti-goat (Jackson Immuno Research, 102649-368) secondary antibodies at 1:500 dilution for two nights at 37°C. Samples were washed 3×1 hour in PBS/TritonX-100 with heparin, then placed through isopropanol/1X PBS dehydration gradient, at 30% for 30 minutes, 50% for 45 minutes, 70% for 30 minutes, and twice in 100% for 20 minutes. Samples were cleared in ethyl cinnamate (Thermo Fisher Scientific, cat# A12906.36) and then mounted with a coverslip’n’slide (NIH 3D 3DPX-009765)^74^ and imaged immediately.

#### Tissue homogenization and RNA extraction

Uterine horns stored in RNA*later* (ThermoFisher, cat# AM7020) were added into 350 µl lysis buffer (1% β-mercaptoethanol in 1x RLT lysis buffer, provided in Qiagen Kit, cat #74106) in 3 mm zirconium bead-containing 2mL tubes (Thermo Fisher, cat# 50-193-1132) and homogenized with a BeadBug 6 (Benchmark Scientific, cat# Z742684) using the following cycles: 60s @ 4350 rpm, 30s rest, 3 cycles total. If tissue was not entirely homogenized, sequential cycles were completed as necessary. Lysate was centrifuged for 2 min @ 22640 rcf (15,000 rpm) through a QIAshredder column (Qiagen, cat #79656) to complete homogenization and RNA was extracted using a RNeasy Plus Kit (Qiagen, cat #74106, #79256) per manufacturer protocol. After RNA was extracted, RNA concentration and quality was measured using a Nanodrop 2000 spectrophotometer. RNA samples were kept at-80°C for long-term storage.

#### Bulk RNA-sequencing analysis and visualization

Downstream analysis of bulk RNA-sequencing was performed in Rstudio. Files containing all differentially expressed genes (DEGs) for estrus and diestrus controls, cHets, and cKOs were read into Rstudio with readxl^63^. *Volcano plots.* The number of upregulated and downregulated genes were counted based on log2 fold change greater than 1.0 or less than-1.0 and p-value being less than 0.05. Plots were created with EnhancedVolcano^69^. *Heatmaps*. Files containing all DEGs were filtered using dplyr^67^ for the gene biotypes “protein coding” and “long noncoding RNAs” (lncRNA) DEGs with a p-value less than 0.01. Each sample’s fragment per kilobase per million (FPKM) values were selected and plotted. Gene IDs were selected for row names, and data was converted to a numeric matrix before calculating the z-score for each DEG with scale function. Final heatmaps were plotted with pheatmap^70^. *GO term.* clusterProlifer^71^, org.Mm.eg.db from Bioconductor^72^, writexl^64^, and ggplot2^73^ packages were utilized. enrichGO from clusterProfiler and *mus musculus* genome data was run on all files. The barplot^60^ function was used to create bar plots of GO terms, showing 15 categories in order of p-value. Writexl was used to save the DEGs from the resulting GO term analysis. Artificial Intelligence was used for help on coding.

#### Bulk RNA-seq data availability

Sequencing data deposited in the Gene Expression Omnibus (GEO) will be publicly accessible after peer review. Please contact authors for earlier release of this dataset.

#### qPCR

*cDNA synthesis*. cDNA was synthesized using manufacturers’ protocol. 1 µg of RNA was added per reaction, and all kit reagents (1x iScript Reaction Mix, 1x iScript Reverse transcriptase, nuclease-free H_2_O) (BioRad, cat #1708890) were combined on ice. cDNA synthesis thermocycler conditions were the following: priming: 5min @ 25 C, reverse transcription: 20 min @ 46°C, reverse transcription inactivation: 1 min @ 95°C. Stock cDNA was stored at 4°C for short-term (up to 1 month) and-80°C for long-term storage. *RT-qPCR.* Prior to qPCR, 5 ng/ul cDNA samples (containing 1x yellow buffer) and 8 µM forward/reverse primer mixes (**Table 3**) were prepared in nuclease-free H_2_O. Novel primers were validated using BLAT^75^ to ensure a single hit within the genome, and melt curves were manually assessed to confirm a single peak. qPCR samples (1x SYBR Green master mix, 400 nM forward/reverse primers) were prepared using the PowerTrack SYBR Green Master Mix kit (ThermoFisher, cat #A46012). Per reaction, the following amounts were added to a 384-well plate per reaction: 0.5 ng/µL DNA, 1x yellow sample buffer, 1x SYBR Green Master mix, 400 nM forward/reverse primers. Each biological sample was completed in triplicate. Negative controls (no DNA added) were included for each gene to ensure lack of DNA contamination. The following cycling conditions were used for qPCR: hold (1x): (+1.6°C/S), 2 min @ 50°C, (+1.6°C/S), 10 min @ 95°C; PCR (40x): 15s @ 95°C, (-1.6°C/S), 60s @ 60°C; melt curve (continuous): (+1.6°C/S), 15s @ 95°C, (-1.6°C/S), 60 s @ 60°C, (+0.075°C/S) 1s @ 95°C. *Quantification/Analysis.* Melt curves were analyzed to validate primer specificity of each gene. 2^-**ΔΔCq**^ analysis was used to quantify relative gene expression Briefly, 3 housekeeping gene raw Cq technical replicate averages and geomeans were calculated for each biological replicate. The following calculations were completed for each biological sample per gene: ΔCq = (Average Cq)-(HK geomean), ΔΔCq = (ΔCq)-(ctrl average ΔCq), 2^-ΔΔCq^ = (2^-ΔΔCq^). Control diestrus was the control (normalization) group. *Hprt*, *Dolk*, and *Pda12* were used as housekeeping genes. Cq values were excluded if they were >37 or undetermined.

**Table 3.**
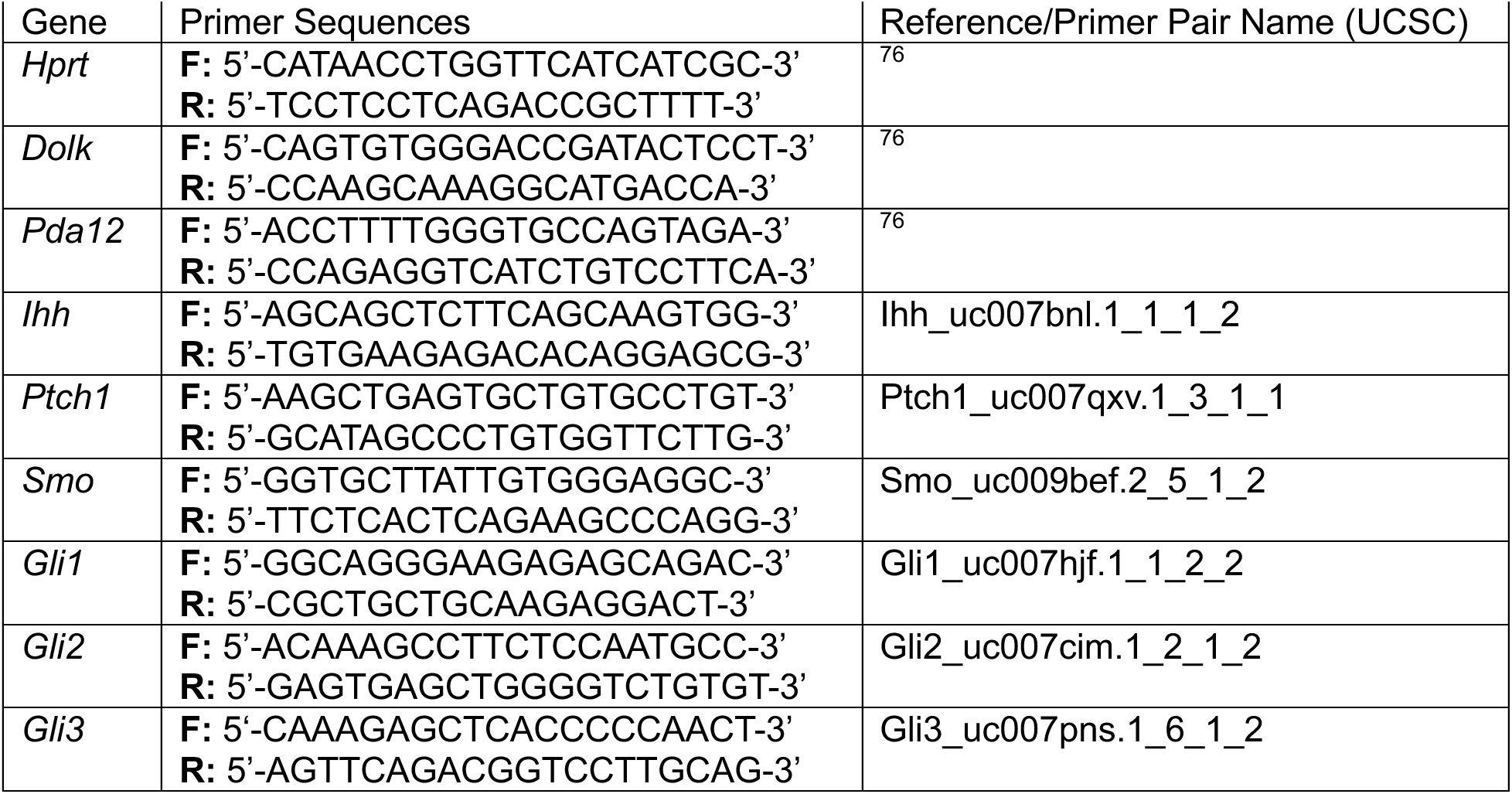

## Supplementary Figures

**Supplemental Figure 1:**
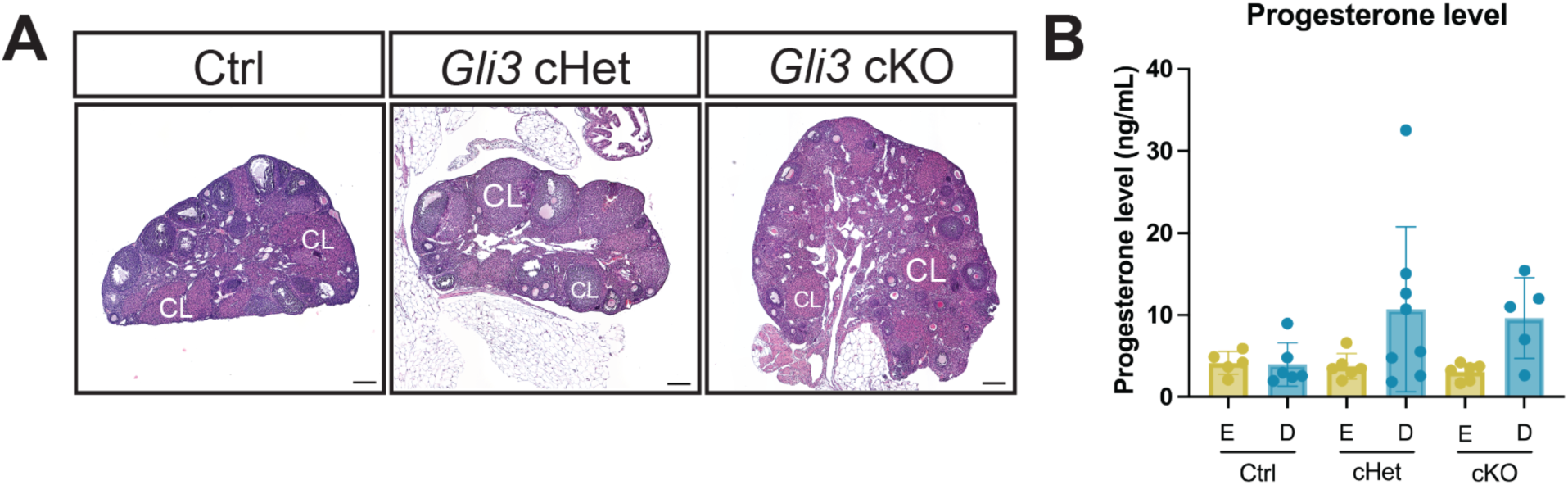
Gli3 is not required for progesterone secretion. (A-C) Hematoxylin and eosin staining of Gli3 control, cHet, and cKO ovaries from estrous cycle checked animals. Imaged with a 10X objective on a Leica DM6b brightfield microscope. Scale bars are 200µm. CL = corpus luteum. **(D)** The average level of progesterone (ng/mL) for estrous cycle checked animals, where each dot is a biological sample.

**Supplemental Figure 2:**
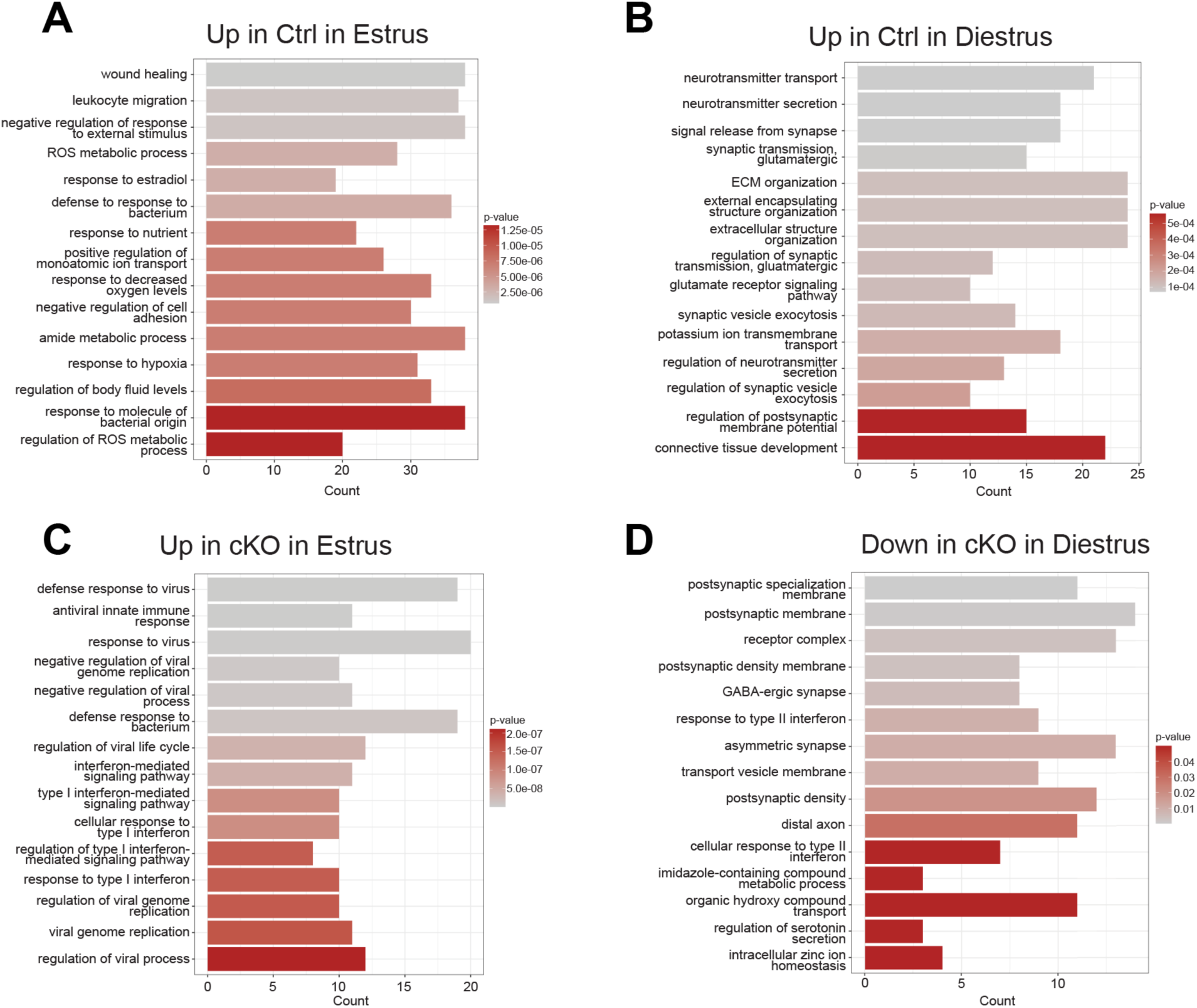
GO term analysis in the Gli3 animals across the estrous cycle. (A-D) GO terms significantly associated with controls at estrus (A) and diestrus (B), as well as Gli3 cKOs at estrus (C) and diestrus (D).

**Supplemental Figure 3:**
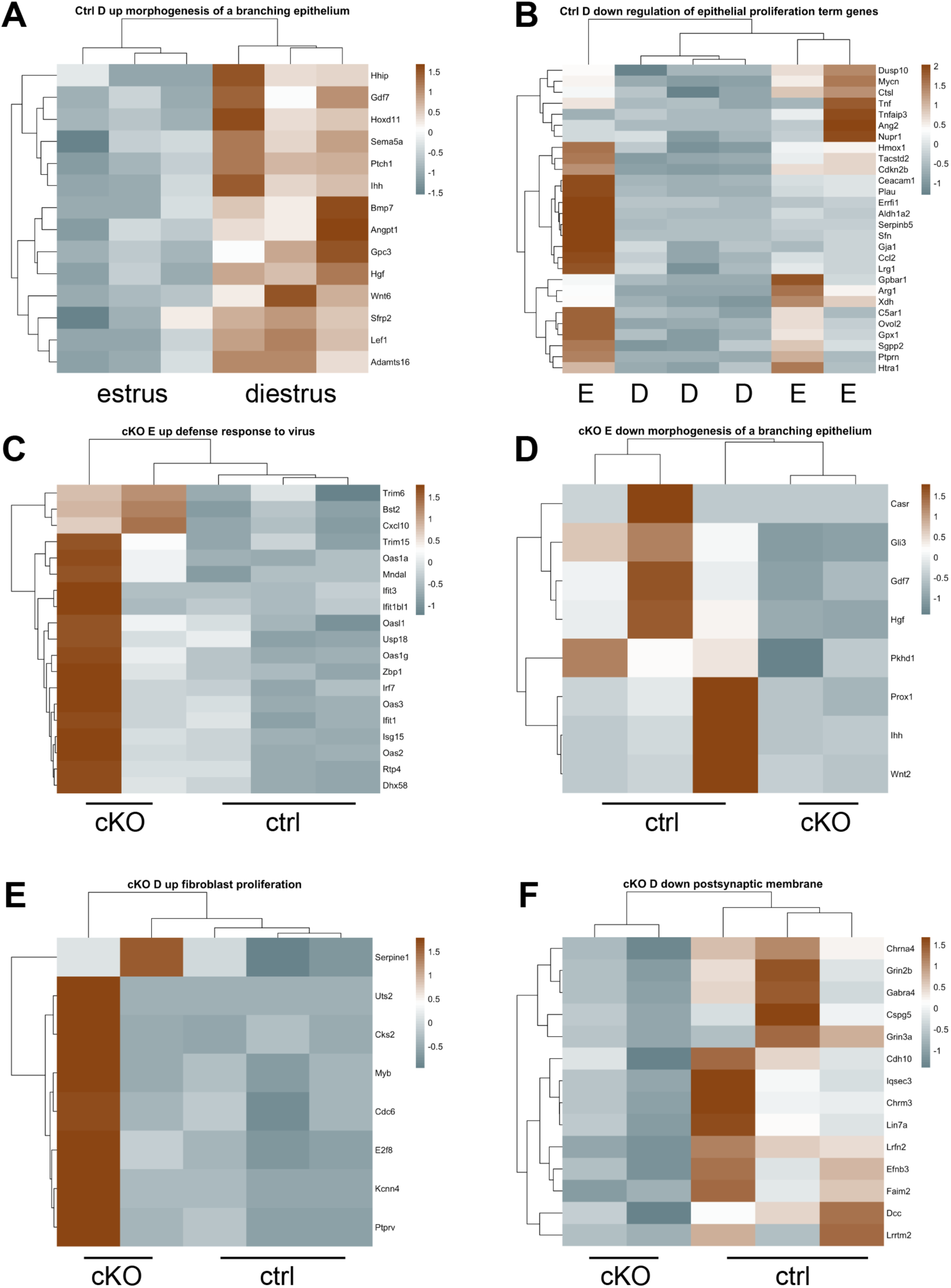
Differentially expressed genes associated with loss of Gli3. (A-F) Heatmaps displaying representative genes associated with different GO terms, including morphogenesis of branched epithelium at diestrus (A), epithelial proliferation at diestrus (B), viral defense response in the cKO at estrus (C), epithelial morphogenesis at in the cKO at estrus (D), fibroblast proliferation in the cKO at diestrus (E), and postsynaptic membrane in the cKO at diestrus (F).

**Supplemental Figure 4:**
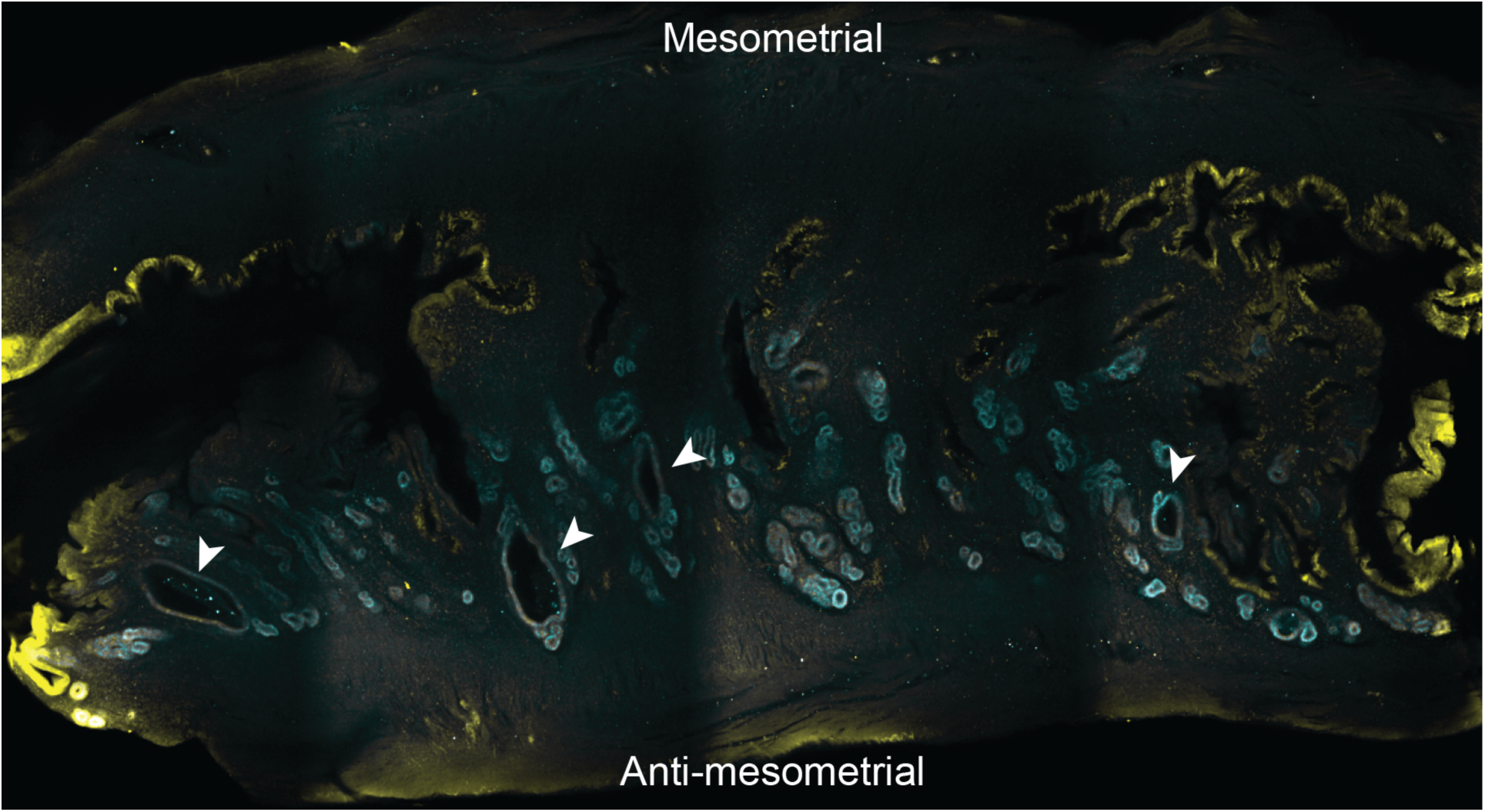
Large glands appear throughout the Gli3 cKO uterus. Whole mount tissue clearing and immunofluorescent staining of Gli3 cKO estrus uterus with Foxa2 (cyan) and E-cadherin (yellow). Arrowheads indicate large glands. Imaged with 10X objective on Andor Dragonfly 200 spinning disk confocal microscope.

**Supplemental Figure 5:**
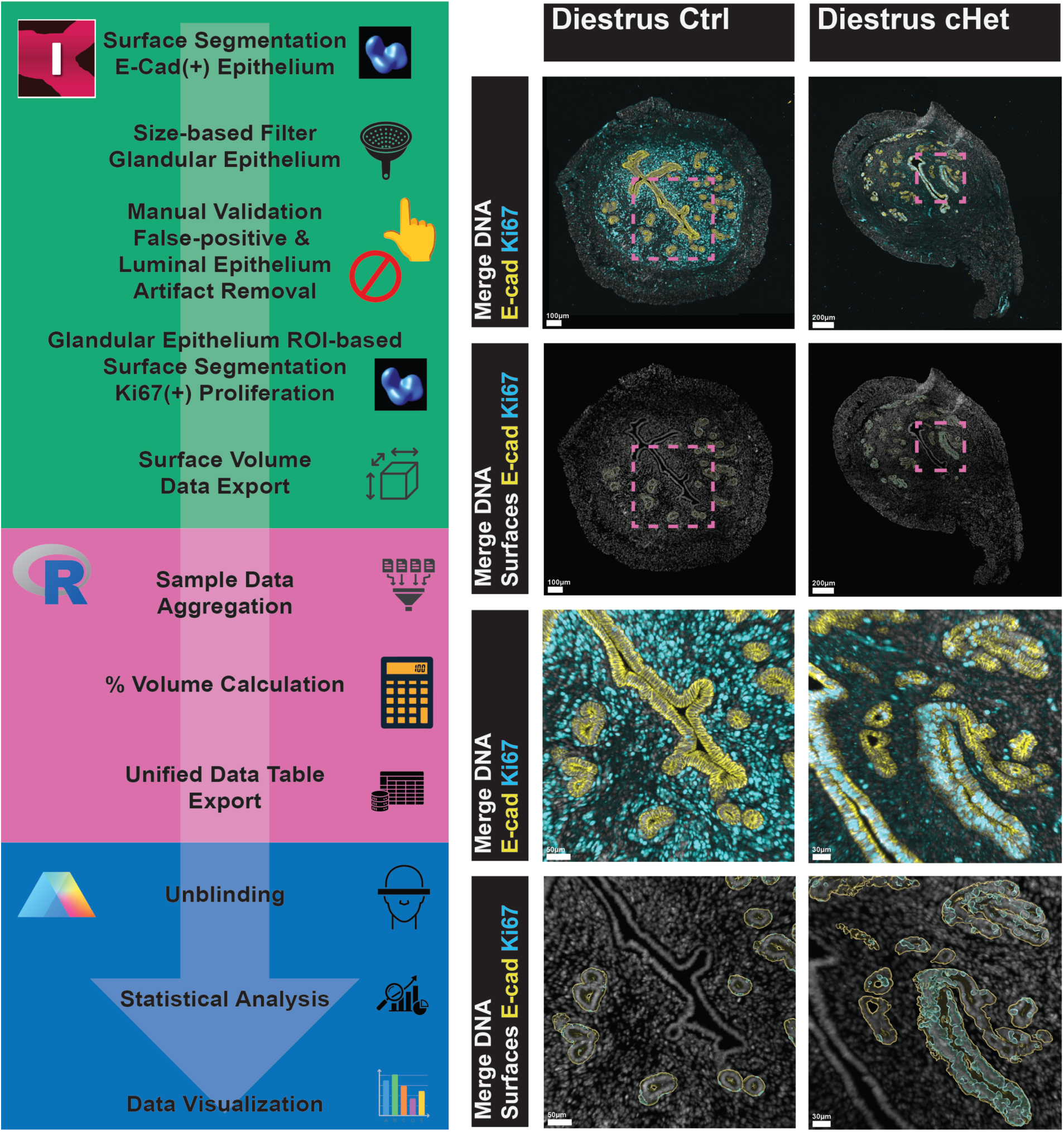
Imaris-based WorkFlow to quantify gland epithelium proliferation.

## References

1. Mouhayar Y, Franasiak JM, Sharara FI. Obstetrical complications of thin endometrium in assisted reproductive technologies: a systematic review. J Assist Reprod Genet. 2019;36(4):607–611. doi:10.1007/s10815-019-01407-y

2. Saad-Naguib MH, Kenfack Y, Sherman LS, Chafitz OB, Morelli SS. Impaired receptivity of thin endometrium: therapeutic potential of mesenchymal stem cells. Front Endocrinol. 2024;14:1268990. doi:10.3389/fendo.2023.1268990

3. Ang CJ, Skokan TD, McKinley KL. Mechanisms of Regeneration and Fibrosis in the Endometrium. Annu Rev Cell Dev Biol. 2023;39:197–221. doi:10.1146/annurev-cellbio-011723-021442

4. Garry R, Hart R, Karthigasu KA, Burke C. A re-appraisal of the morphological changes within the endometrium during menstruation: a hysteroscopic, histological and scanning electron microscopic study. Hum Reprod. 2009;24(6):1393–1401. doi:10.1093/humrep/dep036

5. Critchley HOD, Babayev E, Bulun SE, et al. Menstruation: science and society. Am J Obstet Gynecol. 2020;223(5):624–664. doi:10.1016/j.ajog.2020.06.004

6. Maenhoudt N, De Moor A, Vankelecom H. Modeling Endometrium Biology and Disease. J Pers Med. 2022;12(7):1048. doi:10.3390/jpm12071048

7. Holden EC, Dodge LE, Sneeringer R, Moragianni VA, Penzias AS, Hacker MR. Thicker endometrial linings are associated with better IVF outcomes: a cohort of 6331 women. Hum Fertil. 2018;21(4):288–293. doi:10.1080/14647273.2017.1334130

8. Brodeur TY, Hanson B, Maredia NN, et al. Increasing Endometrial Thickness Beyond 8 mm Does Not Alter Clinical Pregnancy Rate After Single Euploid Embryo Transfer. Reprod Sci. 2024;31(4):1045–1052. doi:10.1007/s43032-023-01385-8

9. Wood GA, Fata JE, Watson KLM, Khokha R. Circulating hormones and estrous stage predict cellular and stromal remodeling in murine uterus. Reproduction. 2007;133(5):1035–1044. doi:10.1530/REP-06-0302

10. Roberson EC, Tran NK, Godambe AN, et al. Hedgehog signaling is required for endometrial remodeling and myometrial homeostasis in the cycling mouse uterus. iScience. 2023;26(10):107993. doi:10.1016/j.isci.2023.107993

11. Lee K, Jeong J, Kwak I, et al. Indian hedgehog is a major mediator of progesterone signaling in the mouse uterus. Nat Genet. 2006;38(10):1204–1209. doi:10.1038/ng1874

12. Franco HL, Lee KY, Broaddus RR, et al. Ablation of Indian Hedgehog in the Murine Uterus Results in Decreased Cell Cycle Progression, Aberrant Epidermal Growth Factor Signaling, and Increased Estrogen Signaling1. Biol Reprod. 2010;82(4):783–790. doi:10.1095/biolreprod.109.080259

13. Harman RM, Cowan RG, Ren Y, Quirk SM. Reduced signaling through the hedgehog pathway in the uterine stroma causes deferred implantation and embryonic loss. Reprod Camb Engl. 2011;141(5):665–674. doi:10.1530/REP-10-0468

14. Wang X, Li X, Wang T, et al. SOX17 regulates uterine epithelial–stromal cross-talk acting via a distal enhancer upstream of Ihh. Nat Commun. 2018;9(1):4421. doi:10.1038/s41467-018-06652-w

15. Wei Q, Levens ED, Stefansson L, Nieman LK. Indian Hedgehog and Its Targets in Human Endometrium: Menstrual Cycle Expression and Response to CDB-2914. J Clin Endocrinol Metab. 2010;95(12):5330–5337. doi:10.1210/jc.2010-0637

16. Lv H, Zhao G, Jiang P, et al. Deciphering the endometrial niche of human thin endometrium at single-cell resolution. Proc Natl Acad Sci U S A. 2022;119(8):e2115912119. doi:10.1073/pnas.2115912119

17. Choi Y, Kim HR, Lim EJ, et al. Integrative Analyses of Uterine Transcriptome and MicroRNAome Reveal Compromised LIF-STAT3 Signaling and Progesterone Response in the Endometrium of Patients with Recurrent/Repeated Implantation Failure (RIF). PloS One. 2016;11(6):e0157696. doi:10.1371/journal.pone.0157696

18. Luo L, Luo M, Ning D, Chen X, Zheng Q, Cao Q. Mechanism of action of IHH in ameliorating thin endometrium. J Reprod Dev. 2025;71(3):161–167. doi:10.1262/jrd.2024-096

19. Wong SY, Reiter JF. The primary cilium at the crossroads of mammalian hedgehog signaling. Curr Top Dev Biol. 2008;85:225–260. doi:10.1016/S0070-2153(08)00809-0

20. Marigo V, Johnson RL, Vortkamp A, Tabin CJ. Sonic hedgehog differentially regulates expression of GLI and GLI3 during limb development. Dev Biol. 1996;180(1):273–283. doi:10.1006/dbio.1996.0300

21. Litingtung Y, Dahn RD, Li Y, Fallon JF, Chiang C. Shh and Gli3 are dispensable for limb skeleton formation but regulate digit number and identity. Nature. 2002;418(6901):979–983. doi:10.1038/nature01033

22. Hui C chung, Angers S. Gli Proteins in Development and Disease. Annu Rev Cell Dev Biol. 2011;27(Volume 27, 2011):513-537. doi:10.1146/annurev-cellbio-092910-154048

23. He F, Akbari P, Mo R, et al. Adult Gli2+/–;Gli3Δ699/+ Male and Female Mice Display a Spectrum of Genital Malformation. PLOS ONE. 2016;11(11):e0165958. doi:10.1371/journal.pone.0165958

24. Blaess S, Stephen D, Joyner AL. Gli3 coordinates three-dimensional patterning and growth of the tectum and cerebellum by integrating Shh and Fgf8 signaling. Development. 2008;135(12):2093–2103. doi:10.1242/dev.015990

25. Yang CF, Chiang MC, Gray DC, et al. Sexually dimorphic neurons in the ventromedial hypothalamus govern mating in both sexes and aggression in males. Cell. 2013;153(4):896–909. doi:10.1016/j.cell.2013.04.017

26. Simon L, Spiewak KA, Ekman GC, et al. Stromal Progesterone Receptors Mediate Induction of Indian Hedgehog (IHH) in Uterine Epithelium and Its Downstream Targets in Uterine Stroma. Endocrinology. 2009;150(8):3871–3876. doi:10.1210/en.2008-1691

27. Matsumoto H, Zhao X, Das SK, Hogan BLM, Dey SK. Indian hedgehog as a progesterone-responsive factor mediating epithelial-mesenchymal interactions in the mouse uterus. Dev Biol. 2002;245(2):280–290. doi:10.1006/dbio.2002.0645

28. The Gene Ontology Consortium, Aleksander SA, Balhoff JP, et al. The Gene Ontology knowledgebase in 2026. Nucleic Acids Res. 2026;54(D1):D1779-D1792. doi:10.1093/nar/gkaf1292

29. Ashburner M, Ball CA, Blake JA, et al. Gene Ontology: tool for the unification of biology. Nat Genet. 2000;25(1):25–29. doi:10.1038/75556

30. Wood GA, Fata JE, Watson KLM, Khokha R. Circulating hormones and estrous stage predict cellular and stromal remodeling in murine uterus. Reprod Camb Engl. 2007;133(5):1035–1044. doi:10.1530/REP-06-0302

31. Jin S. Bipotent stem cells support the cyclical regeneration of endometrial epithelium of the murine uterus. Proc Natl Acad Sci U S A. 2019;116(14):6848–6857. doi:10.1073/pnas.1814597116

32. Syed SM, Kumar M, Ghosh A, et al. Endometrial Axin2+ Cells Drive Epithelial Homeostasis, Regeneration, and Cancer following Oncogenic Transformation. Cell Stem Cell. 2020;26(1):64–80.e13. doi:10.1016/j.stem.2019.11.012

33. Silverberg SG. Problems in the Differential Diagnosis of Endometrial Hyperplasia and Carcinoma. Mod Pathol. 2000;13(3):309–327. doi:10.1038/modpathol.3880053

34. Gonzalez G, Mehra S, Wang Y, Akiyama H, Behringer RR. Sox9 overexpression in uterine epithelia induces endometrial gland hyperplasia. Differ Res Biol Divers. 2016;92(4):204–215. doi:10.1016/j.diff.2016.05.006

35. Jeong JW, Kwak I, Lee KY, et al. Foxa2 Is Essential for Mouse Endometrial Gland Development and Fertility. Biol Reprod. 2010;83(3):396–403. doi:10.1095/biolreprod.109.083154

36. Kelleher AM, Peng W, Pru JK, Pru CA, DeMayo FJ, Spencer TE. Forkhead box a2 (FOXA2) is essential for uterine function and fertility. Proc Natl Acad Sci U S A. 2017;114(6):E1018–E1026. doi:10.1073/pnas.1618433114

37. Jia Z, Li B, Matsuo M, et al. Foxa2-dependent uterine glandular cell differentiation is essential for successful implantation. Nat Commun. 2025;16(1):2465. doi:10.1038/s41467-025-57848-w

38. Johnson DR. Extra-toes: anew mutant gene causing multiple abnormalities in the mouse. J Embryol Exp Morphol. 1967;17(3):543–581.

39. Franco HL, Lee KY, Broaddus RR, et al. Ablation of Indian Hedgehog in the Murine Uterus Results in Decreased Cell Cycle Progression, Aberrant Epidermal Growth Factor Signaling, and Increased Estrogen Signaling1. Biol Reprod. 2010;82(4):783–790. doi:10.1095/biolreprod.109.080259

40. Franco HL, Lee KY, Rubel CA, et al. Constitutive Activation of Smoothened Leads to Female Infertility and Altered Uterine Differentiation in the Mouse. Biol Reprod. 2010;82(5):991–999. doi:10.1095/biolreprod.109.081513

41. Takamoto N, Zhao B, Tsai SY, DeMayo FJ. Identification of Indian Hedgehog as a Progesterone-Responsive Gene in the Murine Uterus. Mol Endocrinol. 2002;16(10):2338–2348. doi:10.1210/me.2001-0154

42. McDermott A, Gustafsson M, Elsam T, Hui CC, Emerson CP, Borycki AG. Gli2 and Gli3 have redundant and context-dependent function in skeletal muscle formation. Dev Camb Engl. 2005;132(2):345–357. doi:10.1242/dev.01537

43. Lex RK, Vokes SA. Timing is everything: Transcriptional repression is not the default mode for regulating Hedgehog signaling. BioEssays News Rev Mol Cell Dev Biol. 2022;44(12):e2200139. doi:10.1002/bies.202200139

44. Lex RK, Zhou W, Ji Z, et al. GLI transcriptional repression is inert prior to Hedgehog pathway activation. Nat Commun. 2022;13(1):808. doi:10.1038/s41467-022-28485-4

45. Zhou Y, Yan H, Liu W, et al. A multi-tissue transcriptomic landscape of female mice in estrus and diestrus provides clues for precision medicine. Front Cell Dev Biol. 2022;10:983712. doi:10.3389/fcell.2022.983712

46. Zhang ET, Wells KL, Bergman AJ, Ryan EE, Steinmetz LM, Baker JC. Uterine injury during diestrus leads to placental and embryonic defects in future pregnancies in mice†. Biol Reprod. 2024;110(4):819–833. doi:10.1093/biolre/ioae001

47. Cooke JP. Inflammation and Its Role in Regeneration and Repair. Circ Res. 2019;124(8):1166–1168. doi:10.1161/CIRCRESAHA.118.314669

48. Rathod S, Shanoo A, Acharya N. Endometriosis: A Comprehensive Exploration of Inflammatory Mechanisms and Fertility Implications. Cureus. 2024;16(8):e66128. doi:10.7759/cureus.66128

49. He Y, Guo Q, Cheng Y, et al. Abnormal activation of the sonic hedgehog signaling pathway in endometriosis and its diagnostic potency. Fertil Steril. 2018;110(1):128–136.e2. doi:10.1016/j.fertnstert.2018.02.138

50. Lees CW, Zacharias WJ, Tremelling M, et al. Analysis of germline GLI1 variation implicates hedgehog signalling in the regulation of intestinal inflammatory pathways. PLoS Med. 2008;5(12):e239. doi:10.1371/journal.pmed.0050239

51. Vue Z, Gonzalez G, Stewart CA, Mehra S, Behringer RR. Volumetric imaging of the developing prepubertal mouse uterine epithelium using light sheet microscopy. Mol Reprod Dev. 2018;85(5):397–405. doi:10.1002/mrd.22973

52. Vue Z, Behringer RR. Epithelial morphogenesis in the perinatal mouse uterus. Dev Dyn Off Publ Am Assoc Anat. 2020;249(11):1377–1386. doi:10.1002/dvdy.234

53. Arora R, Fries A, Oelerich K, et al. Insights from imaging the implanting embryo and the uterine environment in three dimensions. Dev Camb Engl. 2016;143(24):4749–4754. doi:10.1242/dev.144386

54. Granger K, Fitch S, Shen M, et al. Murine uterine gland branching is necessary for gland function in implantation. Mol Hum Reprod. 2024;30(6):gaae020. doi:10.1093/molehr/gaae020

55. Lan Q, Trela E, Lindström R, et al. Mesenchyme instructs growth while epithelium directs branching in the mouse mammary gland. eLife. 2024;13:e93326. doi:10.7554/eLife.93326

56. Byers SL, Wiles MV, Dunn SL, Taft RA. Mouse Estrous Cycle Identification Tool and Images. PLOS ONE. 2012;7(4):e35538. doi:10.1371/journal.pone.0035538

57. Folts L, Martinez AS, McKey J. Tissue clearing and imaging approaches for in toto analysis of the reproductive system†. Biol Reprod. 2024;110(6):1041–1054. doi:10.1093/biolre/ioad182

58. Schindelin J, Arganda-Carreras I, Frise E, et al. Fiji: an open-source platform for biological-image analysis. Nat Methods. 2012;9(7):676–682. doi:10.1038/nmeth.2019

59. Bankhead P, Loughrey MB, Fernández JA, et al. QuPath: Open source software for digital pathology image analysis. Sci Rep. 2017;7(1):16878. doi:10.1038/s41598-017-17204-5

60. R Core Team. R: A Language and Environment for Statistical Computing. Published online 2021.

61. R. Studio Team. RStudio: Integrated Development Environment for R. Published online 2020.

62. Wickham H, Averick M, Bryan J, et al. Welcome to the tidyverse. J Open Source Softw. 2019;4(43):1686–1686. doi:10.21105/joss.01686

63. Wickham H, Bryan J, Posit, et al. readxl: Read Excel Files. Published online March 7, 2025. Accessed April 16, 2026. https://cloud.r-project.org/web/packages/readxl/index.html

64. Ooms J, McNamara J. writexl: Export Data Frames to Excel “xlsx” Format. Published online April 15, 2025. https://cloud.r-project.org/web/packages/writexl/index.html

65. Schauberger P, Walker A. openxlsx: Read, Write and Edit xlsx Files. Published online 2021. https://cran.r-project.org/web/packages/openxlsx/index.html

66. Wickham H, Henry L, Posit Software, PBC. purrr: Functional Programming Tools. Published online April 10, 2026. https://cloud.r-project.org/web/packages/purrr/index.html

67. Wickham H, François R, Henry L, Müller K. dplyr: A Grammar of Data Manipulation. Published online 2021.

68. Wickham H. The Split-Apply-Combine Strategy for Data Analysis. J Stat Softw. 2011;40(1):1–29.

69. Blighe K. kevinblighe/EnhancedVolcano. Published online May 15, 2026. Accessed May 19, 2026. https://github.com/kevinblighe/EnhancedVolcano

70. Kolde R. raivokolde/pheatmap. Published online April 28, 2026. Accessed May 19, 2026. https://github.com/raivokolde/pheatmap

71. Yu G, Wang LG, Han Y, He QY. clusterProfiler: an R Package for Comparing Biological Themes Among Gene Clusters. OMICS J Integr Biol. 2012;16(5):284–287. doi:10.1089/omi.2011.0118

72. Access the Bioconductor Project Package Repository. Accessed May 19, 2026. https://bioconductor.github.io/BiocManager/

73. Create Elegant Data Visualisations Using the Grammar of Graphics. Accessed May 19, 2026. https://ggplot2.tidyverse.org/

74. McKey J, Bunce C, Batchvarov IS, Ornitz DM, Capel B. Neural crest-derived neurons invade the ovary but not the testis during mouse gonad development. Proc Natl Acad Sci U S A. 2019;116(12):5570–5575. doi:10.1073/pnas.1814930116

75. Kent WJ. BLAT--the BLAST-like alignment tool. Genome Res. 2002;12(4):656–664. doi:10.1101/gr.229202

76. Kopinke D, Roberson EC, Reiter JF. Ciliary Hedgehog Signaling Restricts Injury-Induced Adipogenesis. Cell. 2017;170(2):340–351.e12. doi:10.1016/j.cell.2017.06.035

